# Transcription amplification by nuclear speckle association

**DOI:** 10.1101/604298

**Authors:** Jiah Kim, Nimish Khanna, Andrew S. Belmont

**Author notes:** Correspondence to: Andrew S. Belmont.

## Abstract

A significant fraction of active chromosome regions and genes reproducibly position near nuclear speckles, but the functional significance of this positioning is unknown. Here we show that Hsp70 BAC transgenes and endogenous genes turn on 2-4 mins after heat shock irrespective of their distance to nuclear speckles. However, we observe 12-56-fold and 3-7-fold higher transcription levels for speckle-associated Hsp70 transgenes and endogenous genes, respectively, after 1-2 hrs heat shock. Several fold higher transcription levels for several genes flanking the Hsp70 locus also correlate with speckle-association at 37 °C. Live-cell imaging reveals this modulation of Hsp70 transcription temporally correlates with speckle association/disassociation. Our results demonstrate stochastic gene expression dependent on positioning relative to a liquid-droplet nuclear compartment through a “transcriptional amplification” mechanism distinct from transcriptional bursting.

## Introduction

Striking variations in transcriptional activity have been correlated with nuclear compartmentalization. Across multiple species and cell types, lamin-associated domains (LADs), as revealed by DamID, show low gene densities and transcriptional activity (Kind et al., 2013). Similarly, across multiple species and cell types, the radial positioning of gene loci within a cell population stochastically closer to the center of the nucleus is associated with higher transcriptional activity (Kolbl et al., 2012; Takizawa et al., 2008). This stochastic correlation between gene expression and radial positioning may mask a more deterministic relationship between gene expression and gene positioning relative to a specific nuclear body which itself is radially distributed. Nuclear speckles, a RNP-containing, liquid droplet-like nuclear body enriched in both RNA processing and transcription related factors (Lamond and Spector, 2003; Spector and Lamond, 2011), are a prime candidate for such a nuclear body. Nuclear speckles indeed show a radial distribution with decreased numbers near the nuclear periphery and increased concentration towards the nuclear interior. By electron microscopy they appear as interchromatin granule clusters (IGCs)- clusters of ∼20 nm diameter RNPs lying between chromatin regions.

Nuclear speckles were suggested to act as a gene expression “hub” for a subset of genes based on the observation of ∼10/20 highly active genes localizing near the nuclear speckle periphery (Brown et al., 2008; Hall et al., 2006; Shopland et al., 2003). Support for this expression hub model was significantly boosted recently by a new genomic mapping method, TSA-Seq (Chen et al., 2018), which demonstrated that chromosome regions localizing most closely with nuclear speckles correspond largely to the A1 Hi-C subcompartment, one of two major transcriptionally active chromosomal subcompartments, as mapped by Hi-C (Rao et al., 2014). These nuclear speckle-associated chromosome regions were enriched in the most highly expressed genes, house-keeping genes, and genes with low transcriptional pausing. Another new genomic mapping method, SPRITE (Quinodoz et al., 2018), also showed that a large fraction of the genome with high levels of active pol II transcription preferentially positioned near nuclear speckles.

This positioning of a subset of genes near nuclear speckles, however, is only a correlation. Despite this genome-wide demonstration of a subset of active genes positioning deterministically near nuclear speckles, there is no evidence that alleles of endogenous genes actually show different expression levels as a function of speckle proximity. Indeed, the prevailing view has been that nuclear speckles act instead primarily as a storage site for RNA processing factors (Lamond and Spector, 2003).

Previously, we demonstrated an increased speckle-association of BAC transgenes containing the Hsp70 gene locus, including HSPA1A, HSPA1B, and HSPA1L after heat shock (Hu et al., 2009). This increased speckle association was also observed for large, multi-copy insertions of plasmid transgenes containing just the HSPA1A gene and shown to depend on the HSPA1 promoter and proximal promoter sequences rather than the actual transcribed sequences (Hu et al., 2010). Live-cell imaging revealed that the increased speckle association after heat shock for a large, ∼700-copy HSPA1A plasmid transgene array occurred either through nucleation of a new nuclear speckle adjacent to the transgene array, or, more interestingly, through the actin-dependent, long-range directed movement of the transgene array to a preexisting nuclear speckle (Khanna et al., 2014). Strikingly, a significant increase in the MS2-tagged HSPA1A transcript occurred only after but within several minutes after first contact with a nuclear speckle (Khanna et al., 2014).

However, the physiological relevance of this increased transcriptional signal after speckle association of this large plasmid transgene array remained unclear with regard to the actual behavior of the endogenous Hsp70 locus. Cytologically, like other large, heterochromatic plasmid transgene arrays, this HSPA1A transgene array showed an unusually condensed chromatin mass during interphase that was preferentially positioned near the nuclear periphery. Moreover, in contrast to the synchronous induction of transcriptional activation 2-5 mins after heat shock of the endogenous Hsp70 locus, this plasmid transgene arrays showed a highly asynchronous transcriptional activation over 10-30 mins after heat shock (Hu, 2010).

To determine the influence of speckle proximity on transcriptional activation in a more physiological context, we investigated Hsp70 gene activation as a function of speckle association after heat shock at both the endogenous and BAC transgene loci.

## Results and Discussion

First, we identified human haploid Hap1 and Chinese Hamster CHO cell lines in which the endogenous heat shock locus showed significant populations of both speckle-associated (∼90%) and non-speckle associated (∼10%) alleles (> 0.45 µm) (Fig. S1A-D). In contrast, human K562, Tig3, WI-38, and HCT116 showed near 100% pre-positioning adjacent to nuclear speckles (data not shown, (Tasan et al., 2018)). We next established that integrated BAC human Hsp70 transgene in several independently derived CHO cell clones (Hu et al., 2010; Khanna et al., 2014) showed similar gene positioning relative to nuclear speckles before and after heat shock and changes in transcription with speckle association as seen for the endogenous Hsp70 locus in both Hap1 and CHO cells (Fig. S1), suggesting a mechanism that targets both the endogenous Hsp70 locus and BAC Hsp70 transgenes to nuclear speckles. This Hsp70 BAC, with a deletion of the HSPA1A and HSPA1L genes, contains just the single HSPA1B gene within a 172kb human genomic insert (Hu et al., 2009; Khanna et al., 2014). We focused on clone C7 containing 1-3 Hsp70 BAC copies, as estimated by qPCR, integrated at a single chromosomal locus for further investigation.

Hsp70 BAC transgenes in CHO cells showed near identical speckle association behavior and transcriptional induction dynamics after heat shock as the endogenous Hsp70 locus in Hap1 cells (Fig. 1A, Fig S1). The fraction of human haploid Hap1 nuclei containing a positive RNA FISH Hsp70 nascent transcript signal increased from near 0 to 1 between 0-4 mins after heat shock (Fig. 1A, n=110-210, each time point, each replicate). A near identical synchronous induction between 0-4 mins after heat shock was observed for the HSPA1B gene in the Hsp70 BAC transgene (Fig. 1A, n=95-150 each time point, each replicate).

**Figure 1.**
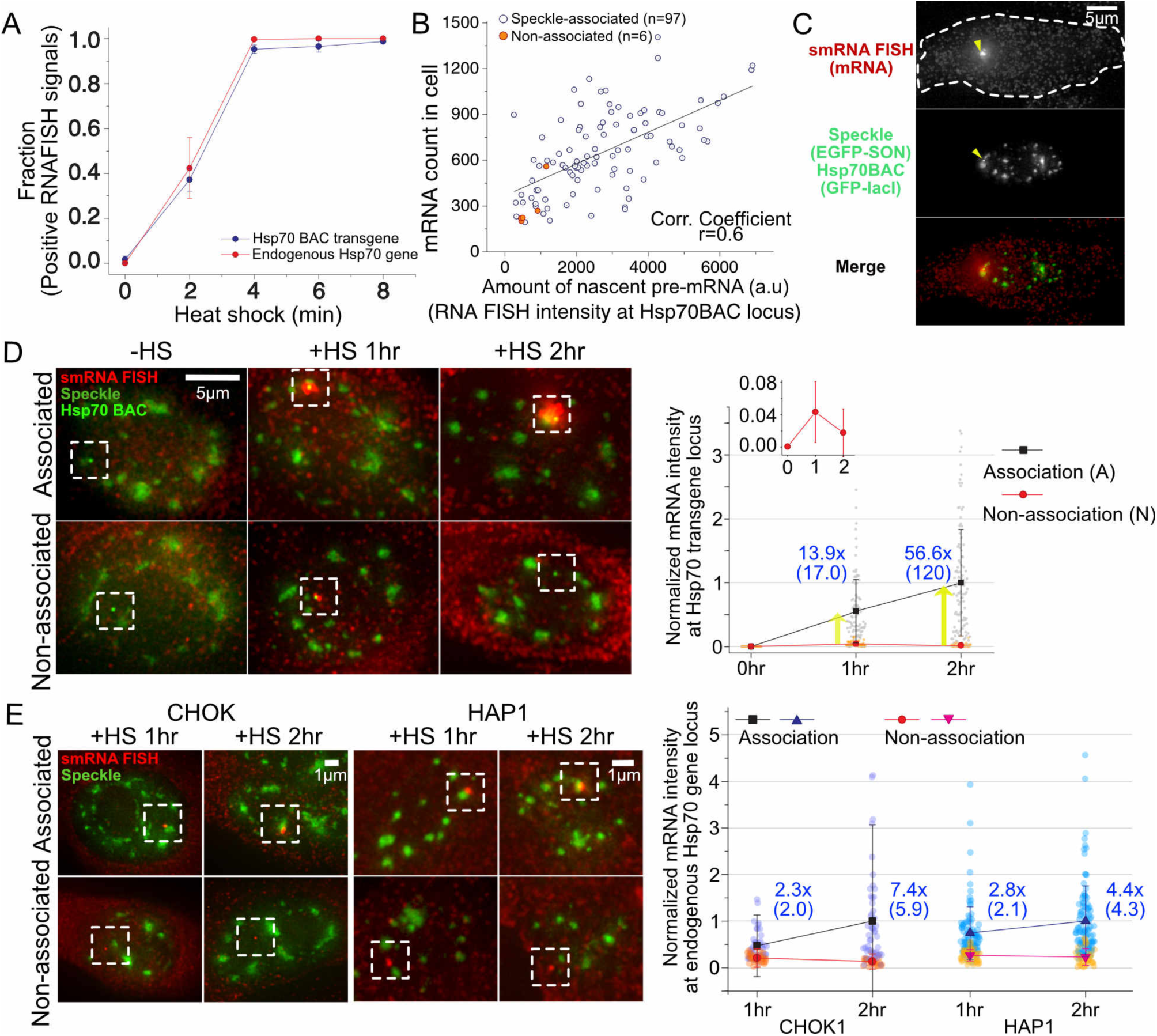
Both Hsp70 BAC transgenes and endogenous genes induce synchronously 2-4 mins after heat shock but show higher transcript levels when associated with nuclear speckles. (A) Transcriptional induction of HSP70 BAC transgene in CHO cells and endogenous Hsp70 genes in HAP1 cells occurs within 2-4 mins after heat shock (SEM, 3 replicates). (B) Scatterplot between levels of nascent pre-mRNA signals versus numbers of mature mRNAs after 15-min heat shock. (C) smRNA FISH image after 15 min heat shock. Nascent pre-mRNAs (arrowhead, top) at Hsp70 BAC transgene (arrowhead, middle). White dashes outline cell border. (D) Higher pre-mRNA levels (boxed regions) for speckle-associated (top) versus non-associated (bottom) BAC transgenes: (Left) Representative images of smRNA FISH versus nuclear speckles and BAC transgene at 0, 1, and 2 hrs after heat shock; (right) mean normalized pre-mRNA intensities at speckle-associated (black, A) or non-associated (red, N) BAC transgenes 0, 1, and 2 hrs after heat shock, with fold differences (blue) of mean (median) for A versus N. (E) Left: Same as in (D) but for endogenous Hsp70 locus in CHO cells (left 2 panels) versus haploid human Hap1 cells (right two panels). Right: Same as in (D), right, for endogenous Hsp70 locus in CHO (left) versus Hap1 cells (right).

We next measured the smRNA FISH signal (nascent pre-mRNA) at both BAC transgenes and endogenous loci when they were speckle-associated versus non-speckle associated. We defined BAC transgenes and endogenous genes as “speckle-associated” if the transgene and/or nascent transcripts positioned within 0.15 µm from the nuclear speckle edge and “non-speckle associated” when the transgene and/or nascent transcripts located further than 0.45 µm from the nuclear speckle edge. Assuming a steady-state between new transcript synthesis and release of transcripts from the transcription site, then the integrated nascent transcript signal should be proportional to the transcription rate. Indeed, the number of dispersed Hsp70 mRNAs 15 mins after heat shock correlates linearly with the nascent RNA signal (Pearson correlation coefficient, R=0.6, Fig. 1B-C).

Despite the near 100% transcriptional induction of alleles 4 mins after heat shock, the actual levels of nascent Hsp70 transcripts increased significantly when the allele was speckle-associated (“A”) versus not-associated (“N”) (Fig. 1D) (n(A/N)= 101/101, 91/40, 107/63 at 0, 1, 2 hrs). At 1 hr after heat shock, we observed a 14-fold higher nascent transcript level for speckle-associated versus non-associated Hsp70 BAC transgenes. This ratio increased to 57-fold by 2 hrs due an ∼2-fold increase in nascent transcripts for speckle-associated transgenes combined with an ∼2-fold decrease in nascent transcripts for non-associated transgenes (Fig. 1D, inset). For the endogenous Hsp70 genes in both CHO and Hap1 cells, we observed an ∼ 3-fold (1 hr) to ∼7-fold (2 hr) increased level of nascent transcripts from speckle-associated versus non-associated alleles after heat shock (Fig. 1E) (CHO: n(A/N)= 46/33, 50/28 at 1 and 2 hrs; HAP1: n(A/N)=102/44, 112/33 at 1 and 2 hrs). Both the endogenous and BAC speckle-associated Hsp70 genes showed roughly a doubling of nascent transcript levels between 1 and 2 hrs after heat shock, but only the BAC transgenes not associated with speckles showed a decrease in transcript levels between 1 and 2 hrs.

The increased transcription of Hsp70 with speckle association raised the question of whether transcription of genes flanking the Hsp70 locus would similarly show increased levels with speckle association. We measured nascent transcripts from three active human genes-VARS, LSM2, and C6orf48-flanking the Hsp70 genes on the BAC transgene (Fig. 2A-C) stably integrated in CHO cells. By smRNA FISH, all three genes showed nearly ∼100% of alleles with nascent transcripts and a 2.5-3-fold increase in nascent transcript levels with speckle association (Fig. 2B-C).

**Figure 2.**
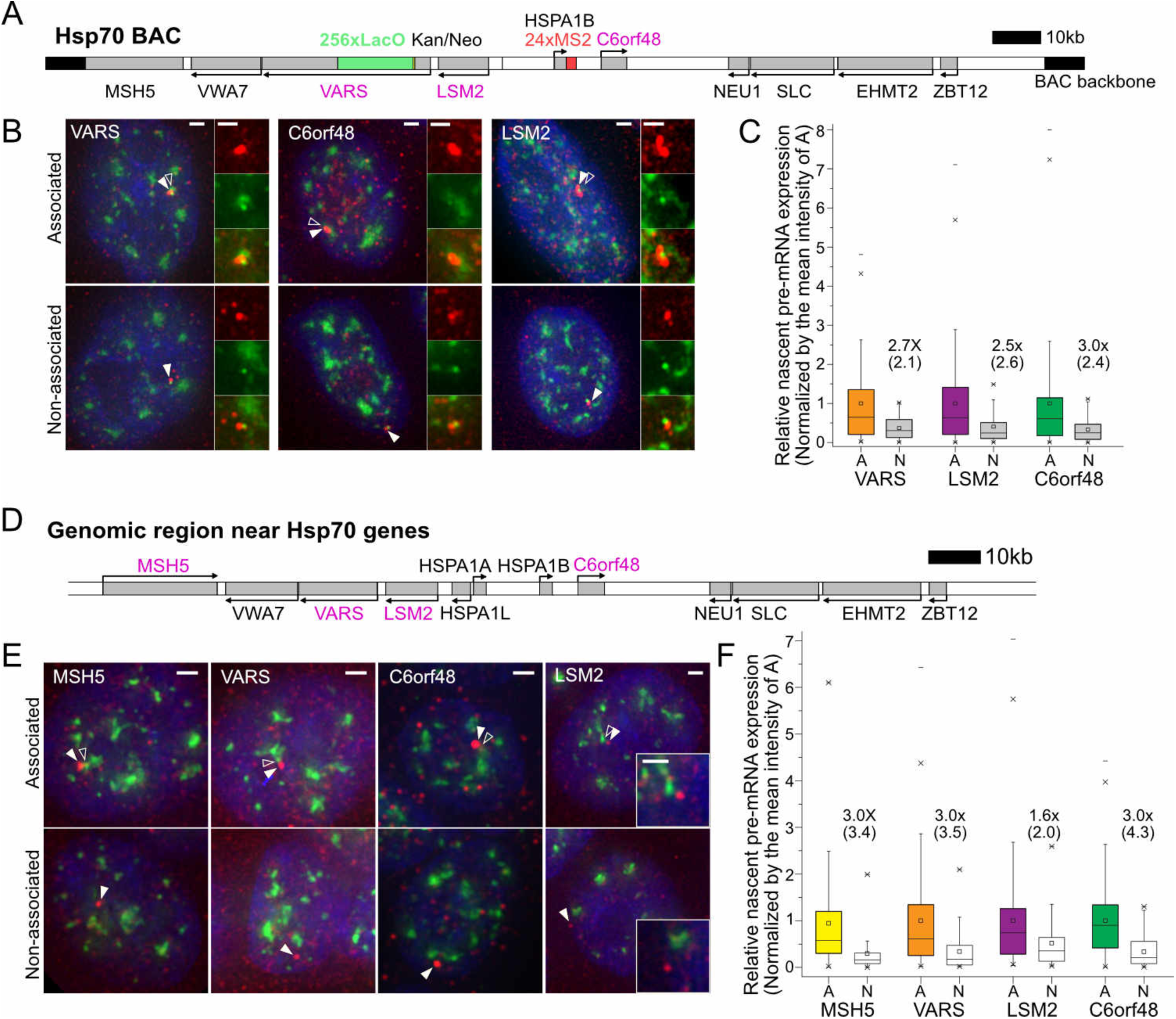
Transcription amplification of speckle-associated genes flanking Hsp70 gene locus at 37 °C. **(A and D)** Probed gene locations (grey boxes, magenta gene names) relative to Hsp70 genes in BAC construct (A) in CHO cells and at endogenous locus (D) in human HAP1 cells. **(B and E)** Representative images of smRNA FISH (red) signals for specific BAC transgene (B) or endogenous gene (E) showing nascent transcripts associated (top) versus non-associated (bottom) with nuclear speckles. White arrowheads-nascent transcripts; Empty arrowheads nuclear speckle. Scale bars, 1 um. **(C and F)** Boxplots showing nascent transcript levels for 3 (C) or 4 (F) genes flanking BAC Hsp70 transgene (C) or endogenous Hsp70 locus (F) as function of speckle association (“A”) or non-association (“N”). Intensities are normalized by the mean intensity at speckle-associated loci: fold differences (x) of the mean (median) between “A” vs. “N” (black). (C) n(A/N)=112/44, 143/37, 185/39 for VARS, LSM2 and C6orf48 BAC transgenes; (D) n(A/N)=168/64, 144/47, 123/52, 203/86 for endogenous MSH5, VARS, LSM2 and C6orf48 genes.

Similarly, for the endogenous human Hsp70 locus we measured 3.0-fold increases for each of the VARS, LSM2, and C6orf48 gene nascent transcript levels and 1.7-fold for the MSH5 gene nascent transcript levels in human Hap1 cells (Fig. 2D-F). In contrast to C6orf48, which showed constant transcriptional activity, MSH5, LSM2, and VARS are bursting genes that show nascent transcripts in only 90±1%, 46±2%, and 70±2% of these Hap1 haploid cells (mean±SEM from 3 independent experiments). However, for each of these three bursting genes, the fraction of cells with visible nascent transcripts associated with nuclear speckles versus visible nascent transcripts not associated with nuclear speckles was comparable to the fraction of gene loci mapped by DNA FISH as adjacent to nuclear speckles versus not adjacent to nuclear speckles (Fig. S1A). Thus, speckle association did not increase the frequency of bursting for these genes, but did enhance their levels of transcription.

To determine the temporal relationship between speckle association and transcription, we used live-cell imaging of the Hsp70 BAC transgenes which contained a 256mer lac operator repeat inserted 29.4 kb upstream of the HSPA1B gene and a 24-mer MS2 repeat inserted into the HSPA1B 3’ UTR (Hu et al., 2009; Khanna et al., 2014). Lac operators were tagged with EGFP-lac repressor, nuclear speckles with EGFP-SON, and the MS2 repeats on transcripts with mCherry-MS2 binding protein (mCherry-MBP).

From 1080 cell movies, we obtained 438 in which the BAC transgene, nuclear speckles, and MS2-tagged transcripts could all be tracked over the entire 25 min observation period. The observed dynamics from each of these 438 cells were then sorted into different categories (Table 1). The three simplest general categories corresponded to cells in which the transgene was always associated with a speckle (Fig. 3A), cells in which the transgene started distant from a speckle but then moved to and remained associated with a speckle (Fig. 3B), and cells in which the transgene became associated with a speckle and showed a visible transcription signal, but then moved away from the speckle (Fig. 4A).

**Table 1.** Categories of dynamics observed for 25mins without or with heat shock.

| Categories | 37°C | Heat shock |
| --- | --- | --- |
| Dynamic range of Hsp70 gene/speckle movement for association/dissociation |  |  |
| $d_{\max} < 0.15 \mu\text{m}$ | 52.7%(79) | 33.3%(146), Fig3A, Video 1 |
| $0.15 \mu\text{m} \leq d_{\max} < 0.45 \mu\text{m}$ | 6.67%(10) | 11.0%(48) |
| $0.45 \mu\text{m} \leq d_{\max}$ | 16.0%(24), | 24.2%(106), Fig3B, 4A&B, Video 2-4, |
| Additional dynamics |  |  |
| Speckle protrusion | 5.33%(8) | 7.53%(33) |
| Speckle formation | 0.00%(0) | 7.99%(35) |
| No association | 0.00%(0) | 1.14%(5), Fig. S3A |
| Gene moving to a different speckle | 6.00%(9) | 6.39%(28) |
| Coordinate movement of speckle&gene | 6.67%(10) | 8.45%(37), Fig. 4C, Video 5 |
| Mitosis | 6.67%(10) | 0.00%(0) |
| Total number of cells | 150 | 438 |
| Faint signal of MCP or BAC or speckles, out of focus | 73 | 642 |

**Figure 3.**
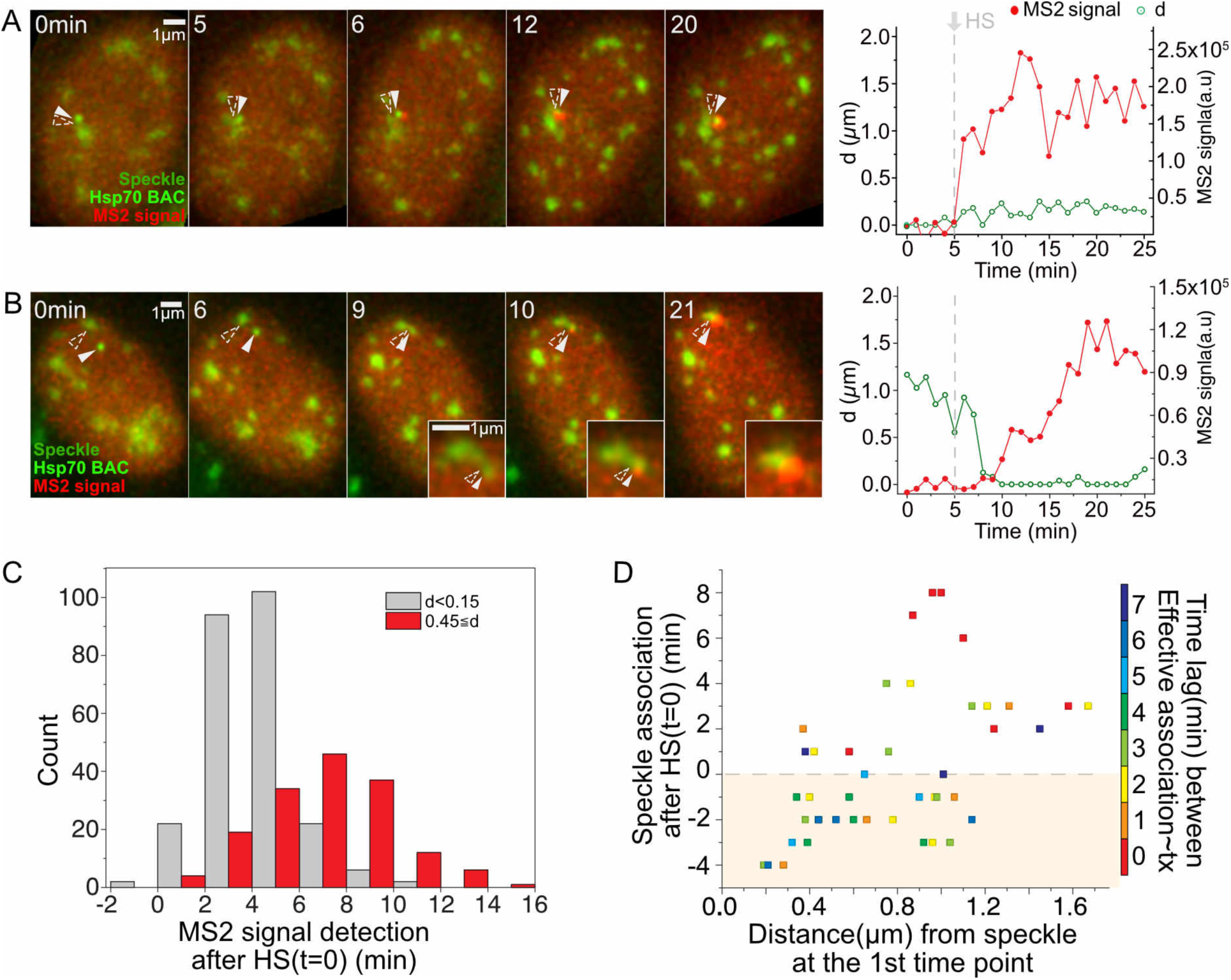
Strict temporal correlation between speckle association and HSPA1B transcriptional amplification. (A & B) (Left panels) Transgene location (solid arrowheads), nuclear speckles (open arrowheads), and MS2 signal versus time (min) after start of observation (Heating on at 1 min, stable heat shock temperature (HS-T) reached at 5 min). (Right panels) Distance (d) of Hsp70 transgene/nascent transcript from closest speckle (green), and nascent pre-mRNA level (red). (A) Transgene associated with speckle throughout HS. MS2 signal appears 1 min after reaching HS-T at 5 mins (Video 1). (B) Transgene initially unassociated with speckle. MS2 signal appears at 10mins, ∼5 mins after reaching HS-T and ∼1 min after moving to and contacting speckle (Video 2). (C) Histogram showing time of MS2 signal appearance after reaching HS-T for transgenes initially speckle-associated (grey) versus not speckle-associated (red). MS2 signal delayed ∼3-mins when not associated (mean=6.5min) versus associated (mean=3.6min). (D) Scatterplot showing timing of speckle-transgene association after reaching HS-T versus initial transgene distance to speckle versus time lag (color) between speckle-gene association and appearance of MS2 signal. Time lags: mean= 3.8 mins, typically 0-3mins after reaching HS-T.

**Figure 4.**
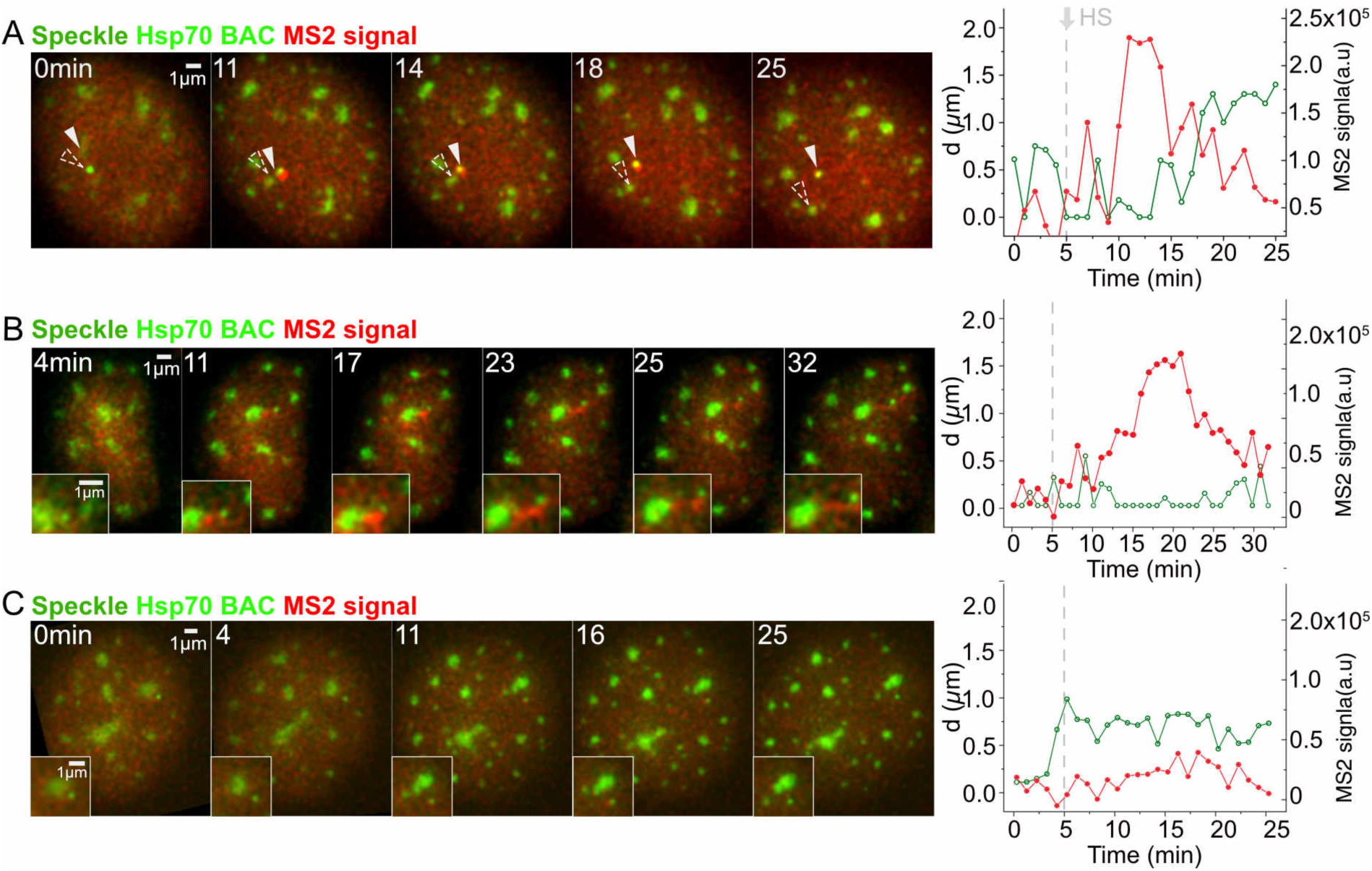
Temporal correlation between speckle dissociation and decrease in HSPA1B transcription. Display and labeling same as in Fig. 3A and B. (A) Hsp70 transgene disassociating from speckle. Nascent transcripts decrease and then disappear after transgene separates from speckle (Video 3). (B) Hsp70 transgene disassociating from speckle. Nascent transcripts accumulate in elongated connection between speckle and transgene after their separation (Video 4). (C) No speckle association of Hsp70 transgene after HS. MS2 signal transiently rises slightly above background but only for single time points (e.g. 16 min).

In the first category (146/438 cells), the BAC transgene remained localized within 0.15 µm from the nuclear speckle during the entire observation period (Fig. 3A). Nascent transcripts became visible above the diffuse background of the MS2-binding protein typically between 2-4 mins after the temperature reached 42°C for heat shock (Fig. 3C, grey bars), and increased gradually afterwards (Video 1). In the second category (41/438 cells), the transgene-speckle distance at some point exceeded 0.45 µm but then the transgene became stably associated with a nuclear speckle and subsequently a MS2 signal appeared (Fig. 3B, Video 2). On average these cells showed an ∼3 min delay in the appearance of a visible MS2 signal relative to cells in the first category (Fig. 3C), due largely to the extra time required for the transgene to move to the speckle. If speckle contact occurred after the temperature had already reached 42°C, a visible MS2 signal typically appeared above background 0-2 mins after contact (Fig. 3B&D). The time lag was longer when speckle association occurred before the temperature reached 42°C (Fig. 3D). Details varied among examples (Table 1); usually the transgene moved to the speckle but occasionally a speckle moved toward the activated transgene (data now shown).

In the third category of speckle movements (28/438 cells), the transgene associated with a nuclear speckle and produced a visible MS2 nascent transcript signal but then moved away from the speckle. Significantly, a decrease or disappearance of the MS2 signal followed this transgene-speckle separation (Fig. 4A, Video 3). Once transgenes moved further than 0.5-1 µm from a speckle, transcripts decayed within 1-2 mins. However, when transgenes moved smaller distances away from the speckle, a low level of transcription was maintained longer rather than a rapid and complete decay of the MS2 signal. When the MS2-transcript level was low, we observed loss of the transcript signal right after transgene-speckle separation, while when transcription levels were higher a complete loss of transcript signal appeared to require a larger separation. Indeed, we could sometimes observe creation of a connecting “bridge” of MS2-tagged transcripts lying between the transgene and speckle periphery. We also could observe deformation of the speckle shape toward the transcripts (Fig. 4B, Video 4). These transcript bridges sometimes elongated as the transgene moved away from the speckle until the transgene had moved far enough away to break contact. Once the transgene dissociated with speckle, the nascent transcript signal did not increase without new speckle association (Fig. 4C).

More complicated and/or rarer classifications shed additional light on the functional significance of nuclear speckle association to HSPA1B transcription (Table 1, Fig. S2). In 5/438 cells, although the transgene did not associate with any speckles, we observed small transient increases for no longer than one time point of the MS2-tagged nascent transcript signal above the MS2-binding protein background (Fig. S3A). In all other cases, appearance of a MS2 signal above background was coupled to nuclear speckle association, which in contrast yielded substantially higher frequency and longer duration increased transcript signals. Additional dynamics include speckle protrusion to a transgene (7.5%), nucleation of a new speckle (8%), movement of the transgene from one speckle to another (6.4%). A physical connection between the BAC transgene, its nascent transcripts, and the associated nuclear speckle is demonstrated by their coordinated movements (8.5%, Fig. S3B, Video 5).

Interestingly, at both heat shock and normal temperatures, we observed long-range transgene movements relative to nuclear speckles (Table 1, Fig. S2). These movements included repetitive oscillations in which transgenes moved large distances away from speckles and then back to the same speckle or sometimes to a different speckle (data not shown). Based on our smRNA FISH results, we anticipate that such oscillations should produce significant variations in the transcriptional levels of the four genes flanking the Hsp70 locus showed transcriptional amplification associated with speckle association as transcription levels of Hsp70 transgenes increase/decrease depending on speckle association/dissociation.

In summary, here we demonstrated a new phenomenon of “transcriptional amplification” for Hsp70 genes and 4 genes flanking the Hsp70 locus, whereby association with nuclear speckles is associated with a several fold boost in transcription. This transcriptional amplification phenomenon is distinct from the now well-described phenomenon of transcriptional bursting, in which genes pulse on and off, in each case for extended time periods.

Our results parallel our previous findings of the appearance of Hsp70 transcription after speckle association of a very large plasmid transgene array. However, this plasmid array was unusually heterochromatic and its activation dynamics after heat shock were greatly delayed and abnormal as compared to the endogenous heat-shock locus. Our current results now extend this earlier work by placing these observations in a physiological context relevant to gene regulation of endogenous gene loci. Moreover, our results now clearly establish that the increased transcription is not related to control of initiation of transcription but rather transcriptional amplification. The tight temporal correlation between speckle association/disassociation and increased/decreased transcription suggests transcriptional amplification follows contact with the nuclear speckle periphery, through direct contact between transgene and speckle and/or through the bridging of nascent transcripts. An actual physical linkage between transgene and speckle through the bridging nascent transcripts is further suggested by live cell movies showing elongation of the MS2 signal during transgene movement away from speckles (Fig. 4B, Video 4) and coordinated movements of speckle, transcript, and transgene (Fig. S3B, Video 5).

Overall, our results support the concept of nuclear speckles as a gene expression hub capable of increasing the level of transcription of associated genes. This transcriptional amplification function of nuclear speckles adds to previously suggested nuclear speckle functions, including modulating of post-transcriptional processing activities such as splicing and nuclear export(Galganski et al., 2017; Spector and Lamond, 2011). We propose speckle association as a new mechanism of stochastic gene expression for a potentially large subset of active genes. Future work will be aimed at determining the prevalence of transcriptional amplification mediated by nuclear speckle contact and its underlying molecular mechanism.

## Materials and Methods

### Cell culture and establishment of cell lines

CHO cells were grown in Ham’s F12 media (Cell Media Facility, University of Illinois at Urbana-Champaign) with 10% fetal bovine serum (FBS) (Sigma-Aldrich, F2442) at 37°C in a 5% CO_2_ incubator. To generate stable CHO cell lines, we carried out a series of DNA transfections followed by selection of stable colonies with 4 different DNA constructs in the following order: 92G8+3’MS2+GKREP_C26 Hsp70 BAC (G418, 400µg/ml), p3’SS-EGFP-dlacI (hygromycin, 200µg/ml) (Robinett et al., 1996), EGFP-SON-Zeo BAC (zeocin, 200µg/ml), and pUb-MS2bp-mCherry (puromycin, 400µg/ml) (Khanna et al., 2014). Modifications of the original 92G8 Hsp70 BAC (Invitrogen) to 92G8+3’MS2+GKREP_C26 included insertion of a lac operator repeat and Kan/Neo selectable marker cassette (Hu et al., 2009), deletion of 8 kb containing the HSPA1A and HSPA1L genes (Hu et al., 2010), and insertion of MS2 repeats into the 3’ UTR of HSPA1B (Khanna et al., 2014). The original SON BAC (165J2, Invitrogen) was modified by adding GFP to the NH2 terminus of the SON coding region to generate EGFP-SON-Zeo BAC (Khanna et al., 2014). The final CHO cell clone we used for live-cell imaging in these studies was C7MCP. C7MCP cells were maintained in complete media with 200 µg/ml G418, 100 µg/ml hygromycin, 100 µg/ml zeocin, 200 µg/ml puromycin. When cells were seeded on coverslips or glass-bottom dishes 2 days prior to cell imaging, media was changed to complete media without any pH indicator and with no G418, hyrogomycin, zeocin, or puromycin added. Hsp70_5, Hsp70_14, Hsp70_20 CHO stable cell clones, generated previously (Hu et al., 2009), contain the full length Hsp70 BAC, modified with the lac operator repeat and selectable marker, integrated at different insertion sites. The parental CHO DG44 cell line used for the generation of these three clones stably expressed EGFP-Lac repressor. These three clones were grown in Ham’s F12 media with 10% FBS with 100µg/ml hygromycin and 200µg/ml G418.

### Single molecule RNA FISH (smRNA FISH) probe design

smRNA FISH was performed using Stellaris probes and the Stellaris protocol (Biosearch Technologies). Probes were designed with the Stellaris Probe Designer using 1.5∼2kb of the 5’ end of coding sequence. FISH probe sets for each gene consisted of ∼33 20mer DNA oligonucleotides, complementary to the target RNA. Probe sets used were: CHO *Hsp70* (cacgcacgagtaggtggtg, cttccgtgctggaacacg, tggtcgttggcgatgatct, gtcggtgaaggccacgtag, cgtcgaaacacggtgttctg, cgaacttgcggccgatcag, gatctcctcggggtagaag, catcttcgtcagcaccatg, tgatcaccgcgttggtcac, gagtcgttgaagtaggcgg, cgtgggctcgttgatgatc, ccaggtcgaagatgagcac, atggacacgtcgaacgtgc, gaagatgccgtcgtcgatc, cacgaagtggctcaccagc, tggacgacagggtcctctt, cctcgaacagggagtcgat, cgccgtgatggacgtgtaga, ggaacaggtccgagcacag, cttctgcaccttggggatg, gttgaagaagtcctgcagc, gatgctcttgttgaggtcg, ctgcacgttctcagacttg, ttgatgagcgccgtcatca, gagtaggtggtgaaggtct, cgtacacctggatcagcac, gatgccgctgagctcgaag, atcgatgtcgaaggtcacc, tgacgttcaggatgccgtt, taggactcgagcgcgttct, gcgctcttcatgttgaagg, aggagatgacctcctgaca, cacgaactcctccttgtcg, ctaatccacctcctcgatg, acaccgggagagcaagcag, agggctaactaaccctgac), *Hsp70* (gggagtcactctcgaaagac, cacaggttcgctctggaaag, aacgccggaaactcaacacg, cgacaagagctcagtcctt, tgagactgggggctggaaac, tggtcgttggcgatgatctc, ttcgcgtcaaacacggtgtt, ttgtctccgtcgttgatcac, agatctcctcggggtagaat, atctccttcatcttggtcag, ctgcgagtcgttgaagtagg, atgatccgcagcacgttgag, tcaggatggacacgtcgaag, tcgaagatgccgtcgtcgat, ccctcaaacagggagtcgat, ggtgatggacgtgtagaagt, ttcggaacaggtcggagcac, aggaccaggtcgtgaatctg, ttgaagaagtcctgcagcag, ttgatgctcttgttcaggtc, tgcacgttctcggacttgtc, ttgatcagggcagtcatcac, tcgtacacctggatcagcac, cagattgttgtctttcgtca, atgtcgaaggtcacctcgat, tgacgttcaggatgccgttg, tcatgttgaaggcgtaggac, tgtccagaaccttcttcttg, cacgagatgacctcttgaca, ttgtgctcaaactcgtcctt, ctgatgatggggttacacac, actaaagaacaaaggcccct, aagtccttgagtcccaacag, ccatcaggttacaacttaac), *MSH5* (ccacgagcctgcaaaagga, cgctacaggtgggagaacg, cgccttttcagtaacctga, tcacgcgcttatcttcctc, gagtcgtgcacgtcttatg, gaaggaaggggtctgaggg, tcattcctgtcacgcggag, attgtgggaaactccacgc, ggtgaattctcgggtattt, tggagagctgtggacacag, tacttctgctacagggctg, tggcgcgcagaatgcaaag, acgattcacagaggaggcc, aaaagtgagggcggttcgg, catgagcttggaggctctg, cttgggttcgctcctaag, ctggggaagccggaggag, ttctactcccctcagagac, ccatcaactctccattcaa, ccaaccctcttttattcta, gatctgtccagcaaggaag, attccacagcacacacaga, taggcaatgcccaagtatc, gtggagtcactagtatcat, gcatctggcatgaagtgga, gagaagcttgaggctctcg, ggaattcatggttccatcc, catctgcaatcccagagag), *VARS* (acacccctgagcacgacg, cgcggacgcaggacgaga, ccggtctcacgaggaaca, ctttgtgacagggagcgt, tagttccctaagatcgcc, agtcgagcgggcagagac, aatccacctcacagccag, agatggtcagactgggcc, cgcgtgtacgtactggag, tcggcgtctggttggatg, ggagtgtggaaggctctc, aactatccaccatcgcgg, ggggagttcctggggaag, tctcagaggggcagtgtc, ctgtcaggagccgaggac, ctgagggtctgaccaggc, agacacccgagtcccata, ctcacgggagctccttcg, ggtcctatgtttgagtag, gaagactgcgggatcgag, aggtccgaacgaagtgga, gggagacgtagagggtgg, ctggggaaggcatctggg, tagcgagcggctatgagg, gtggctggagacagatgc, gttggacaaggacagccg, cgtgtcggcgtaactgac, cacaggcagctggtatta, cgagcttcggagtcccag), *LSM* (gagcgcaagctgggtagag, caagcgctgacgggcaaag, gcgggaagcgacgcagaaa, caggtctggggaaaccgaa, ggaagacagcagggtgctg, cttttgacgtcacggtacc, ggtacaaaggccagatccc, ctcctcaatgaacctgaga, atatcccatttgttctcag, ttttttccctcatcatgga, tcctcactgaatctctctc, aggtctaacttttccgtct, aagcttgcctggcagagaa, agaaaatatcccacccgca, cccactcctttcaatgaat, ctgtgctctctcagtcgac, aagtcccagagagactctg, aaggctggagcccaaatta, catccctgacagttctcaa, tactggtattgtgaacccc, ggagtgtcctttgacagta, tgtgggaaggagcatggta, gggagagagggggaaaacc, tagttccacgaccacatcc, gtacctacctcaggtcatt, cccaacagacttgttggaa, tgctttgtattgttttcca, aaagaggttccagggccc, gtacatcttccacctcgc, aggcacaaggcatttattt, actgcgcctgacctgtatt), *C6orf48*(gagcccacttcgcaaaaag, aactctcatactgccaacc, cactcaacagtcgggccat, tgcgcaaaggcagcgcaag, caacggtagttcacccaac, agacaccagaaactccagt, gagctaggtcagttccaag, ttgttttgagtcagcaggg, ccattttgcaatcactcgc, atcatcactcccttcctat, cactcaagagttacctggg, tcagagcgctgcggtgatg, aaatgctggaccgaggggg, ccatgaactcgttgagcct, ttacaactcctaacgggga, ggataacacggcgatgaac, cctaatactcacctttact, aggcattcaaaaggctctc, ttgggaacaaagctttccg, atgcccatgaacgaaagct, actcctattttgcagtaga, ggacgttagaaagggagga, agggaagctcttctggaaa, ccaattagggagatctgga, agcatcggagactctagtc, gttccctaaatgagtcaga, cttctggagacccaagtat, tgggcttccagagttcatt, ctctgtgaaggtgcattgt, ccttcactcagattagtgc, gagggggagattccaaacc, gccatacaaagcttctctc)

### Single molecule RNA FISH (smRNA FISH) procedure

Each oligonucleotide contained an amino group at the 3’ end for fluorophore coupling using either Cy5 NHS ester (GE Healthcare, PA15102) or rhodamine NHS ester (ThermoFisher, 46406). NHS esters and probes were incubated overnight in 0.1M sodium bicarbonate solution (pH 8.0) at room temperature, and purified using Bio-Spin P6 columns (Bio-Rad, 7326221) according to the manufacturer’s protocol. Purified, pooled probe concentrations were ∼50-100 µM in Tris buffer, pH 7.4.

Cells were seeded on coverslips (Fisher) 2 days before experiments. For heat shock, a well-plate or dish containing the coverslip and media was sealed using parafilm and incubated in a 42°C-water bath. Cells were fixed using freshly prepared 3.6% paraformaldehyde (PFA, Sigma, P6148-500G) in phosphate-buffered saline (PBS) for 15 mins at room temperature (RT). After washing 5 mins 3x in PBS, cells were permeabilized using 0.5 % Triton X-100 (ThermoFisher, 28314) for 10 mins in DEPC-treated PBS. For DEPC treatment, 1 ml fresh DEPC was added to 1 L PBS or water, incubated more than 15 hrs at RT, and autoclaved for 25 min to inactivate the remaining DEPC. After rinsing 3x in DEPC-treated PBS, the permeabilized cells were equilibrated in wash buffer (10% Formamide (Sigma, F9037), 2x saline-sodium citrate (SSC)) for 30 mins. The cells were incubated in hybridization buffer with smRNA FISH probes (final concentration, ∼300-500nM) for 15 hrs at 37 °C. The hybridization buffer contained 2x SSC, 10% formamide, 10% w/v dextran sulfate (Sigma, D8906), 1 mg/ml E.coli tRNA (Sigma, R8759), 2 mM ribonucleoside vanadyl complex (RVC) (NEB, S1402), 0.02% RNase-free BSA (Ambion, AM2618) in DEPC-treated water. The cells were then washed 30 mins 2x at 37 °C in wash buffer, and mounted in a Mowiol-DABCO anti-fade medium (Ed Harlow, 1988).

smRNA FISH in HAP1 cells was followed by immunostaining against SON. After smRNA FISH, cells were fixed again with 3.6% PFA in DEPC-treated PBS for 15 mins, and washed for 5 mins 3x. Cells were incubated in blocking buffer (0.5% Triton X-100, 1% w/v RNase-free BSA, 20 µM RVC). Cells were incubated with primary antibody against SON (Sigma, HPA 023535) at a 1:300 dilution in DEPC-treated PBS for 1 hr in a humid chamber at RT, washed for 5 mins 3x in DEPC-treated PBS, and incubated with Alexa488-labeled, goat anti-rabbit IgG secondary antibody (Invitrogen, A11008) at a 1:300 dilution in DEPC-treated PBS for 1 hr in a humid chamber at RT. All antibody solutions were supplemented with 0.4U/µl RNase inhibitors (Lucigen, E0126). For smRNA FISH against endogenous transcripts in wild-type CHO cells, immunostaining against SC-35 was done prior to smRNA FISH. We followed a similar immunostaining procedure as described above, but with primary antibody against SC35 (1:300 in PBS, Abcam, ab11826) for 12 hrs at 4°C and with Alexa488-labeled, secondary goat anti-mouse IgG antibody (Invitrogen, A11029) at a 1:300 dilution in DEPC-treated PBS. After washing 5 mins 3x in DEPC-treated PBS, slides were mounted in a Mowiol-DABCO anti-fade medium.

### Microscopy and data analysis of fixed samples

For fixed cells, we used a Personal Delta Vision microscope (GE Healthcare) equipped with a Coolsnap HQ camera and Plan Apo N 60x/1.42 NA oil-immersion objective (Olympus). Sections were spaced every 200 nm in z. Pixel size was 67 nm. 3D z-stacks were processed using the “Enhanced” version of the iterative, nonlinear deconvolution algorithm provided by the Softworx software (GE Healthcare) (Agard et al., 1989), and projected in the x-y plane using a maximum intensity algorithm. The distance between a gene and a nuclear speckle was measured from the maximum intensity projection of the 3D data set: the edge of the speckle was defined as where the nuclear speckle intensity fell to 40% of its intensity maximum, and the distance measured to the center of the BAC transgene or the edge of a FISH signal. Measurements were made manually using the line profile function in ImageJ. smRNA FISH signal intensities over nascent transcripts were measured by manually selecting the nascent RNA FISH area in 2D summed projections of the 3D raw data set, summing the pixel intensities, and then subtracting the background intensity estimated from a same size area adjacent to the nascent transcript signal. Measurements were made using ImageJ. For counting the number of mature mRNA spots, we used the StarSearch software program, as described elsewhere (Levesque et al., 2013; Shaffer et al., 2013).

### Live cell imaging and analysis

Cells were plated on 35mm dishes with a #1 1/2 thickness glass coverslip bottom (MatTek, P35G-1.5-14-C) 48 hrs before imaging. For rapid, wide-field live-cell imaging, we used a GE OMX V4 microscope (GE Healthcare) equipped with a U Plan S-Apo 100×/1.40 NA oil-immersion objective (Olympus), two Evolve EMCCD cameras (Photometrics), a live cell incubator chamber (GE Healthcare) with separate temperature controllers for the objective lens and the incubator heater, and a humidified CO2 supply. MatTek dishes were placed on the microscope and temperatures for both the objective lens and incubator chamber were maintained at 37 °C for ∼1-3 hrs prior to data acquisition. Temperatures were set to 44 °C on both temperature controllers immediately after the 1st time frame was taken, with ramping from 37 °C to 44 °C requiring ∼4 mins. Empirically, this temperature setting of 44°C for the controllers produced a media temperature of 42°C inside the Matek dish and a similar transcriptional induction of the Hsp70 BAC transgene MS2 signal on the microscope as seen off the microscope with a 42 °C heat shock.

3D images (z-spacing=200nm) were acquired once every min, typically using the solid-state illumination with 1% transmittance, 10-15 msec exposure for each z-slice for 477±16 nm excitation (GFP) and 2∼5% transmittance, 10-20 msec exposure for each z-slice for 572±10 nm excitation (mCherry). Typically, 3D stacks from each of 20-25 fields of view were taken during each 1 min time point interval using the point-visiting function of the Softworx image acquisition software, with each 3D data stack acquired over ∼ 1 sec or less. 3D z-stacks were processed using the “Enhanced” version of the iterative, nonlinear deconvolution algorithm provided by the Softworx software (Agard et al., 1989) (GE Healthcare), and projected into the x-y plane using a maximum intensity algorithm. Using a custom Matlab program, each cell in the projected live cell movies (512×512 pixel area per frame) was tracked and cropped into a 256×256 pixel area per frame with the cell translated to the center of the area and saved as a single file. If necessary, a rigid body registration (ImageJ plugin ‘StackReg’) was applied to correct for any x-y nuclear rotation and/or translational displacement between sequential time points. Each of these individual cell movie files was then visually inspected and sorted into different categories of dynamics for further analysis.

**Software:** Custom MATLAB scripts written for data analysis described in are publicly available in Github: DOI:10.5281/zenodo.2559675, URL: https://doi.org/10.5281/zenodo.2559675

### Online Supplemental Material

Fig. S1 provides information about Positioning of Hsp70 BAC transgene or endogenous Hsp70 gene relative to a nuclear speckle before (37 °C) or after heat shock (HS). Fig. S2 provides details about more complicated classifications of relative movements between BAC transgene and nuclear speckle. Fig. S3 shows extra examples of interesting speckle and transgene dynamics. Video 1 is the source of Fig. 3A showing stable speckle association and increase in transcription after heat shock. Video 2 is the source of Fig. 3B showing speckle association after heat shock and delayed increase in transcription. Video 3 is the source of Fig. 4A showing dissociation of transgene from speckle and decrease in transcription. Video 4 is the source of Fig. 4B showing dissociation of transgene from speckle with bridging transcripts between transgene and speckle. Video 5 is the source of Fig. S3 showing coordinate movement of transgene and speckle via physical attachment.

## Acknowledgements

### Funding

ASB acknowledges support from National Institutes of Health R01 grant GM58460 and from its Common Fund 4D Nucleome Program (U54 DK107965).

### Author contributions

J.K. and A.S.B conceived of and designed the study; J.K performed experiments and analyzed data; J.K and A.S.B wrote manuscript; N.K provided a stable cell line. Competing interests: The authors declare that they have no competing interests.

### Data and materials availability

Data and materials will be provided upon request to the corresponding author; some reagents may be subject to Material Transfer Agreements.

**Videos:** Time (min) and scale bar (1 µm) are stamped on each video. Videos represent maximum intensity 2D projection of 3D image stack for each time point. Each image of 3D stack was captured every min. Videos play at 20fps. Video 1-5: Temperature increase begins at 1 min, reaching 42°C at ∼5 mins.

**Video 1.** Appearance of MS2-tagged nascent transcripts (red) without delay after heat shock for BAC transgene (green.) stably associated with nuclear speckle (lighter green) (see also Fig. 3A).

**Video 2.** Appearance of MS2-tagged nascent transcripts (red) is delayed after heat shock, with MS2 signal appearing after non-associated transgene (green) moves to and makes contact with nuclear speckle (lighter green) (see also Fig. 3B).

**Video 3.** Decrease and disappearance of MS2-tagged nascent transcripts (red) after disassociation of transgene (green) from nuclear speckle (lighter green) (see also Fig. 4A).

**Video 4.** Delayed decay of MS2-tagged nascent transcripts (red) after transgene (green) disassociation from nuclear speckle (lighter green). This delay is associated with a nascent transcript accumulation between transgene and speckle that elongates and appears to physically connect the nuclear speckle with the transgene even after the transgene moves away from the speckle (see also Fig. 4B).

**Video 5.** Coordinated movement of transgene (green) and nuclear speckle (light green) suggesting stable physical attachment of transgene with speckle during heat shock (see Fig. S7).

## Supplementary figures

**Figure S1.**
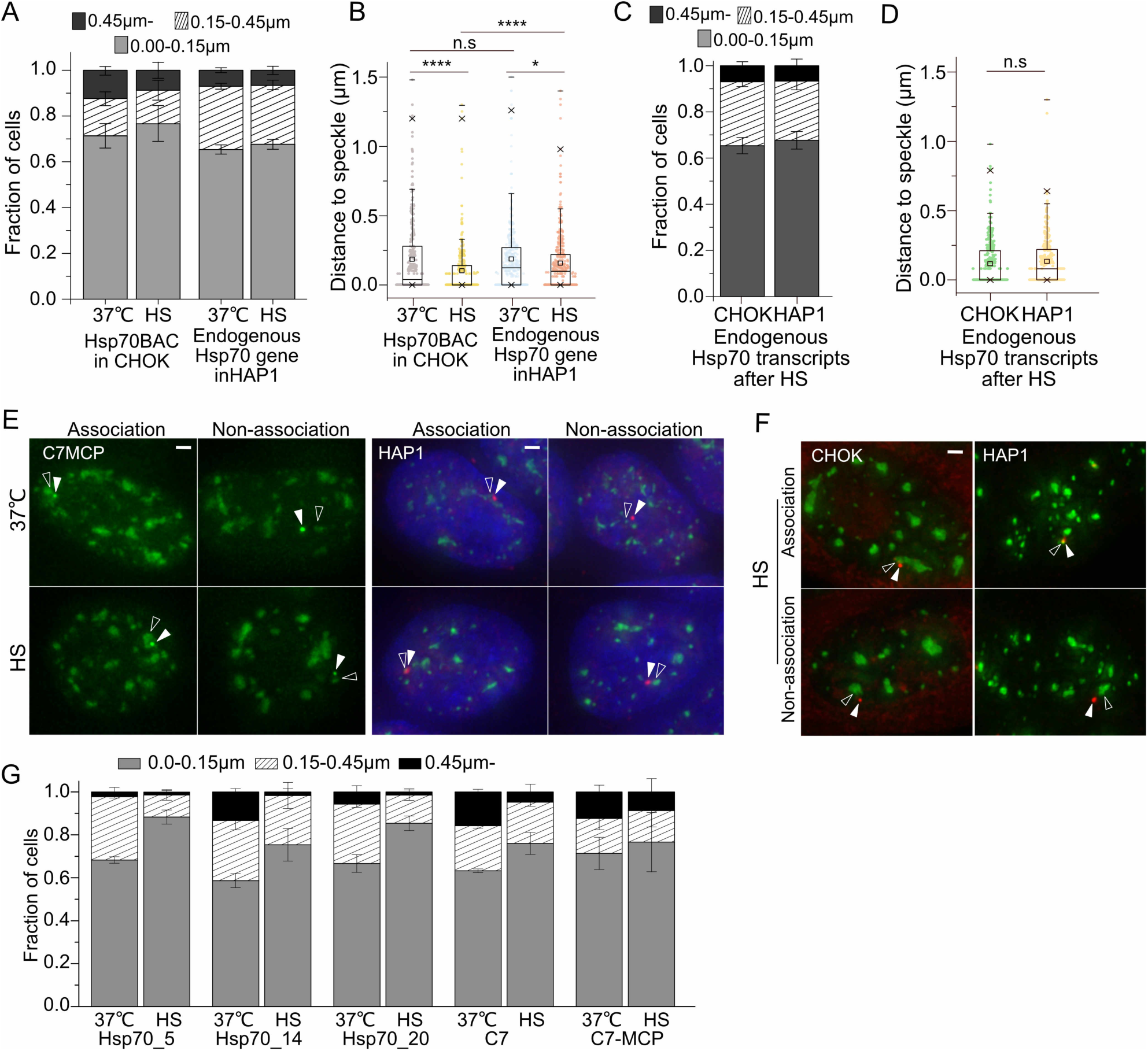
Positioning of Hsp70 BAC transgene or endogenous Hsp70 gene relative to a nuclear speckle before (37 °C) or after heat shock (HS). **(A)** Histograms showing fraction of BAC Hsp70 transgenes (lacO) in CHO cell clone C7MCP or endogenous Hsp70 alleles in HAP1 cells (DNA FISH) at varying distances from the nuclear speckle before and after 30 min HS (mean±SEM, 3 biological replicates, N= 100-170 per replicate). **(B)** Boxplots showing distribution of varying distances from speckle shown in histogram (A). Mean (square inside box), median (line), 25 (bottom) and 75 (top) percentiles; ends of error bars-10 (bottom) and 90 (top) percentiles. *p<0.05, ****p<0.00001, n.s: not significant. Paired Wilcoxon signed rank test was used. **(C)** Histograms showing fractions of RNA FISH signals from the endogenous Hsp70 locus in CHO or HAP1 cells at varying distances from nuclear speckles after 30 min HS (mean±SEM, 3 biological replicates, N= 100-120 per replicate). **(D)** Boxplots showing distribution of varying distances from speckle shown in histogram (C). Box format is same as (B). **(E)** Position of BAC transgene (green, white arrowhead) and nuclear speckle (green, empty arrowhead) (Left panels) or endogenous gene DNA FISH signal (red, white arrowhead) and nuclear speckle (green, empty arrowhead) (Right panels, DAPI staining blue) at 37 °C (top) or after HS (bottom). Scale bars= 1 µm. (F) Position of RNA FISH of endogenous Hsp70 locus transcripts (red, white arrowheads) and nuclear speckle (green, empty arrowheads) after 30 min HS in CHO cells (left) or HAP1 cells (right). Scale bars= 1 µm. **(F)** Positioning of Hsp70 BAC transgenes relative to nearest nuclear speckle before and after 30min HS in several independently derived CHOK cell clones. Fraction of BAC transgenes at different distances relative to nuclear speckle (mean ± SEM, 3 biological replicates, N= 90-150 per replicate).

**Figure S2.**
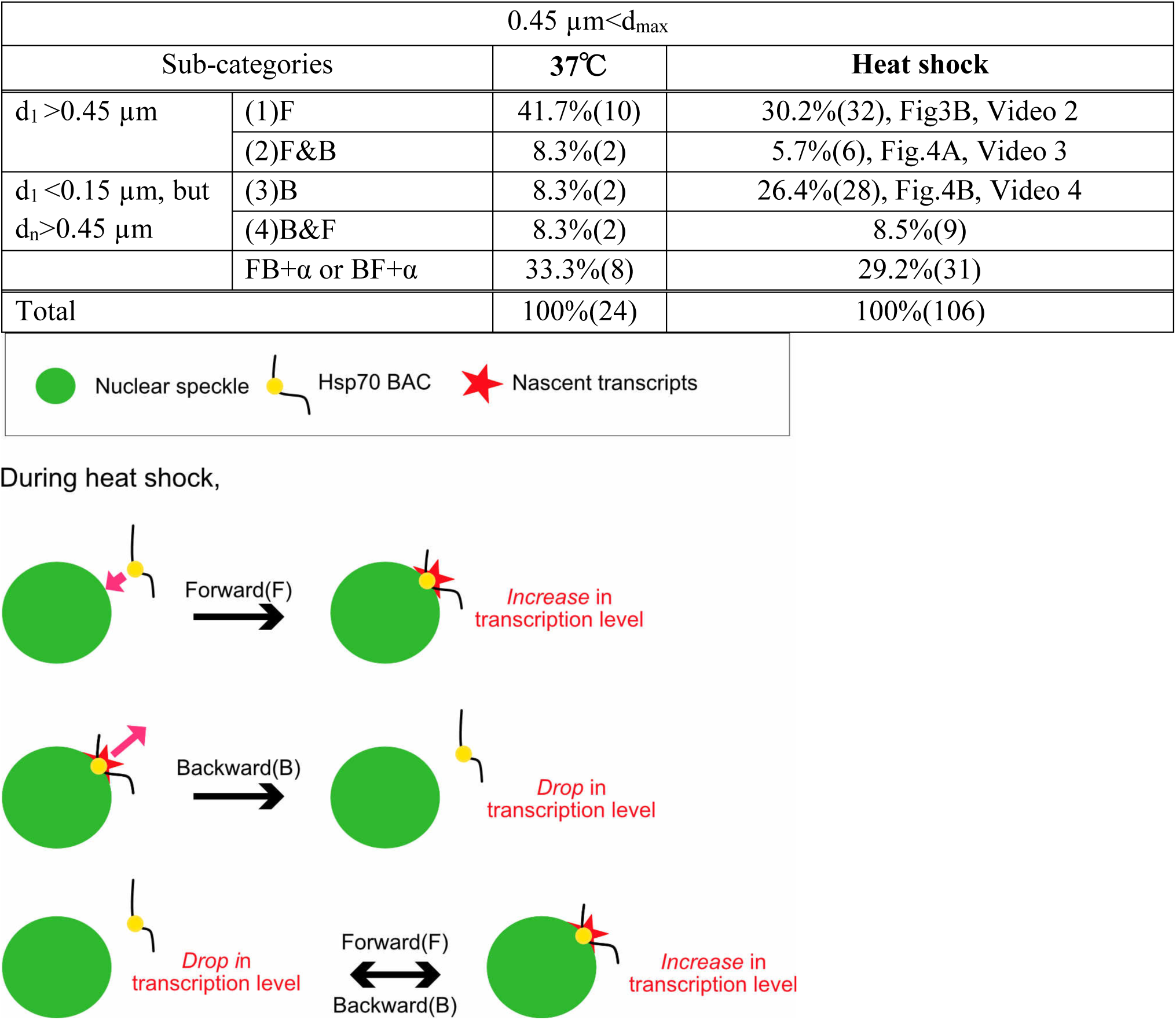
Statistics for more complicated classifications of relative movements between BAC transgene and nuclear speckle. d_max_: Maximum distance between transgene locus and speckle observed during imaging. d_1_: Distance between transgene locus and speckle at the 1^st^ time point of imaging. d_n_: Distance between transgene locus and speckle at any time point during imaging. F: Forward motion to the transgene or speckle toward another. B: Backward motion of the transgene or speckle relative to another. +α: Additional movements.

**Figure S3.**
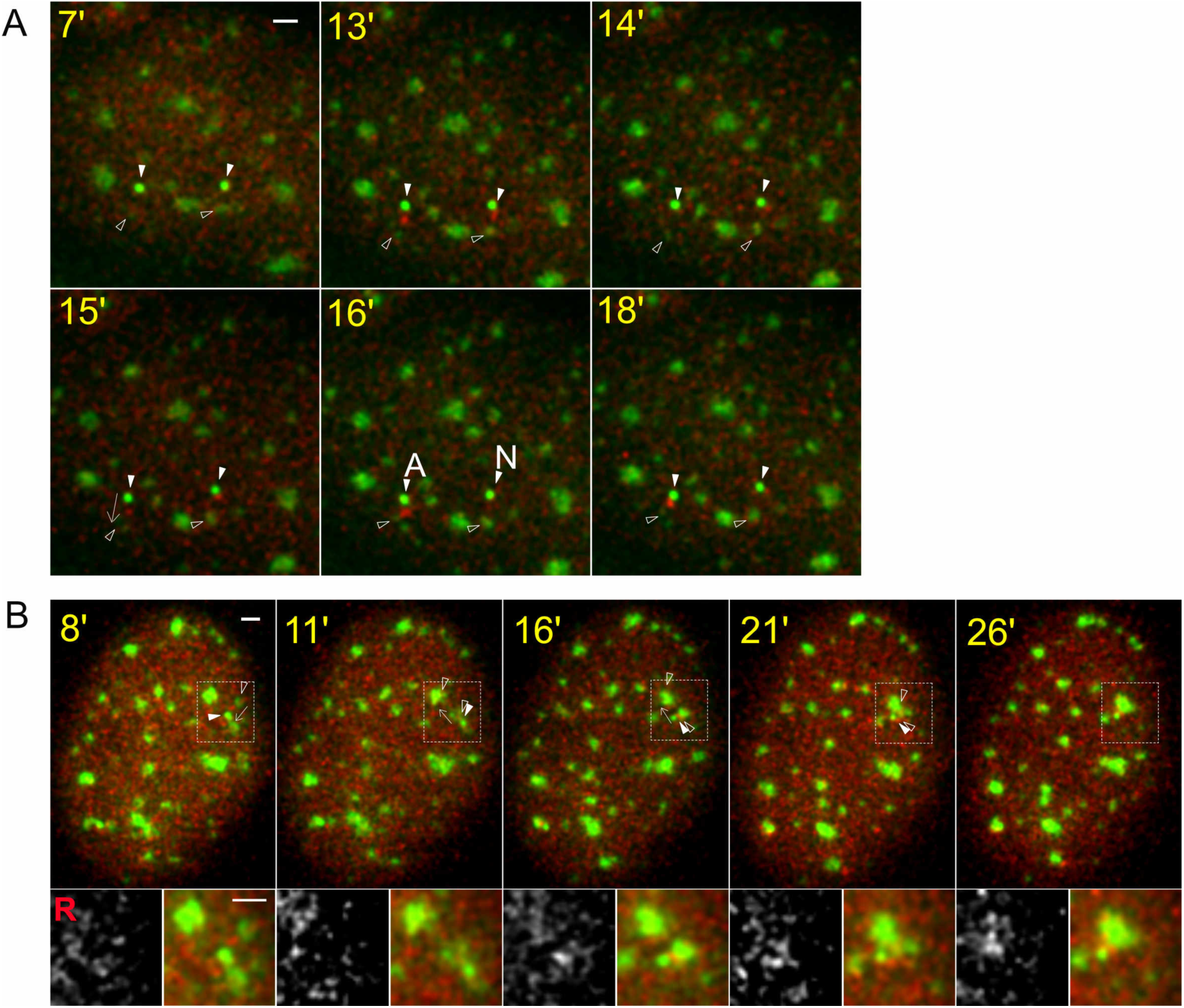
Extra examples of interesting speckle and transgene dynamics. Time stamp during HS (yellow), BAC transgene (white arrowheads, bright green), nuclear speckle (empty arrowheads, lighter green), RNA MS2-tagged transcripts (red, mCherry-MS2 binding protein). Scale bars, 1µm. **(A)** No persisting transcription for non-associating locus. Category: No association, Table 1. Arrow (15 min) shows direction of transgene movement. Transcriptional bursting is observed at both non-associating transgene loci at 13 mins. These bursting signals are observed again at 15 min at both loci. At 16 mins, one transgene (left) associates (“A”) with a speckle and now maintains elevated transcript signal during the rest of observation time, whereas the other transgene which is not associated (“N”) with speckle does not maintain an elevated transcript signal. **(B)** Coordinated movement of speckle and gene. Nuclear speckle and associated BAC transgene move together as a single unit before merging with a different speckle, suggesting a stable attachment of transgene and speckle (Video 5). Category: coordinate movement of speckle & gene, Table 1.

